# Characterization of the intraspecies chimeric mouse brain at embryonic day 12.5

**DOI:** 10.1101/2025.03.31.646380

**Authors:** Phoebe Strell, Madison A. Waldron, Sether Johnson, Anala Shetty, Andrew T. Crane, Clifford J. Steer, Walter C. Low

**Affiliations:** Department of Neurosurgery, Medical School, University of Minnesota, Minneapolis, MN, USA; Stem Cell Institute, Medical School, University of Minnesota, Minneapolis, MN, USA; Department of Veterinary and Biomedical Sciences, University of Minnesota, Saint Paul, MN, USA; Graduate Program in Neuroscience, Medical School, University of Minnesota, Minneapolis, MN, USA; Department of Medicine, University of Minnesota, Minneapolis, MN, USA; Department of Genetics, Cell Biology and Development, University of Minnesota, Minneapolis, MN, USA

## Abstract

Incidence of neurodegenerative diseases such as Alzheimer’s, Parkinson’s, Huntington’s, and amyotrophic lateral sclerosis have increased dramatically as life expectancy at birth has risen year-over-year and the population ages. Neurological changes within the central nervous system, specifically the brain, include cell loss and deterioration that impact motor function, memory, executive function, and mood. Available treatments are limited and often only address symptomatic manifestations of the disease rather than disease progression. Cell transplantation therapy has shown promise for treating neurodegenerative diseases, but a source of autologous cells is required. Blastocyst complementation provides an innovative method for generating those autologous neural cells. By injecting mouse induced pluripotent stem cells (iPSCs) into a wild type (WT) mouse blastocyst, we generated a chimeric mouse brain derived of both donor and host neuronal and non-neuronal cells. An embryonic day 12.5 (E12.5), automated image analysis of mouse-mouse chimeric brains showed the presence of GFP-labeled donor-derived dopaminergic and serotonergic neuronal precursors. GFP-labeled donor-derived cholinergic precursor neurons and non-neuronal microglia-like and macrophage-like cells were also observed using more conventional imaging analysis software. This work demonstrates that the generation of mouse-mouse chimeric neural cells is possible; and that characterization of early neuronal and non-neuronal precursors provides a first step towards utilizing these cells for cell transplantation therapies for neurodegenerative diseases.

## Introduction

Neurodegenerative diseases are incurable, age-dependent, or genetic component disorders that result in cell loss and deterioration in the nervous system^1^. In recent years, the increasing elderly population has driven the prevalence of neurodegenerative diseases such as Alzheimer’s disease, Parkinson’s disease, Huntington’s disease, and amyotrophic lateral sclerosis. These diseases impact memory, thought processing, language, problem-solving, and movement^2–5^. The neuropathology and physiology for these diseases are diverse; however, all of these diseases display a common characteristic in the loss of neurons. In some cases, microglia are implicated in disease pathology and are associated with increased inflammation and immune signaling, compromised blood-brain barrier, stress responses, and more^6,7^. For example, in Alzheimer’s disease there is loss of serotonergic neurons in the dorsal raphe nucleus and cholinergic neurons in the nucleus basalis and medial septal nucleus, while in Parkinson’s disease, there is a loss of dopaminergic neurons in the substantia nigra pars compacta and cholinergic neurons on the nucleus basalis^8^. These brain regions are by no means an exhaustive list of affected areas. The manifestation of symptoms is dependent upon the affected brain regions that undergo neuronal loss^8^. Treatments for neurodegenerative disease are rather limited and mostly include (i) pharmacological agents to slow down the degenerative process or compensate for the neuronal loss; and (ii) non-pharmacological interventions to improve quality of life^9,10^. Therefore, therapeutic approaches to replace the lost neuronal and non-neuronal cells may prove to be more effective. Clinical trials of transplanted dopaminergic neuron precursor cells from human fetal brain tissue have resulted in the amelioration of movement disorders in patients with Parkinson’s disease for over 25 years after transplantation, without the need for levodopa medication^11–13^.

The ethical concerns of using human fetal brain tissue as a source of cells for clinical transplantation, however, have resulted in the exploration of other sources of cells for implantation. The derivation of induced pluripotent stem cells (iPSCs) via reprogramming of adult somatic cells, and the differentiation of these iPSCs into dopamine precursor cells have demonstrated efficacy in transplantation studies in animal models of PD, and are currently being evaluated in a limited number of phase I clinical trials^14–17^. However, studies evaluating the efficacy of differentiated iPSCs compared to primary cells demonstrated limitations. These included different transcriptional expression profiles, limited engraftment of transplanted cells, including short extensions for neurite outgrowth, no synapse formation, requirement for more tyrosine hydroxylase (TH) positive cells in the graft to reverse behaviors, and risk for graft overgrowth in the iPSC derived cells after transplantation^18,19^. The results from these studies suggest that dopamine precursor cells generated from this approach are not completely reprogrammed to function as authentic precursor cells.

Blastocyst complementation is a method to generate authentic exogenic cells, tissues, and complete organs from one species within the body of another that behaves as a biological incubator^20^. A typical of blastocyst complementation approach involves generating a developmental niche by ablating a gene of interest in the early host embryo, for a specific tissue or organ, which is followed by microinjecting donor pluripotent stem cells (PSC) into the genetically modified host blastocyst^21^. The progeny of the donor cells should fill this developmental niche. Past studies have successfully generated intraspecies (e.g., mouse-mouse) and interspecies (e.g., rat-mouse) chimeric liver, forebrain, kidney, pancreas, lung, thyroid, eye, heart, and vascular system via blastocyst complementation^22–28^.

Blastocyst complementation of the nervous system has only begun to be explored in the last decade as a method of repair. *Neurog1*^+/-^-deficient mice complemented with donor mouse PSCs successfully formed donor-derived inner ear sensory neurons, correcting for the typical ear malformations observed in these mice^29^. An additional study used a targeted ablation of dorsal telencephalic progenitors expressing *Emx*1 during development, causing agenesis of the cerebral cortex and hippocampus. Donor embryonic stem cells injected into these genetically engineered blastocysts resulted in morphological and neurological normal brain tissue^30^. More recent studies have begun to explore the brain as an organ that can also be generated through more traditional methods of blastocyst complementation. A recent approach used several gRNAs and CRISPR to generate both intraspecies mouse-mouse and interspecies rat-mouse chimeric animals with chimeric brains^31^. Another group sought to generate wild type rat-mouse chimeras to study the functional circuitry of the rat and mouse neurons^32^. Yet, these studies lack characterization of the specific neural and non-neural cells developing within these chimeric brains. A basic understanding of intraspecies mouse-mouse chimeras can provide insight into donor-derived cellular heterogeneity and the level of mosaicism observed in various brain regions.

Donor cell contribution has become one of the driving outcomes for determining the success of blastocyst complementation or microinjection of donor cells into unmodified blastocysts. Donor cells microinjected into a wild type (WT) blastocysts can contribute and form interspecies chimeras, and mores specifically brain regions^32,33^. As the generation of the nervous system is rather novel among blastocyst complementation studies, a basic understanding of donor cell contribution must be conducted in intraspecies chimeras to provide a foundation for future blastocyst complementation studies. Additionally, to evaluate the use of these cells for future transplantation therapies, understanding what types and what proportions of precursors are present will be helpful to determine if WT chimeras alone can provide a sufficient number of cells for transplantation^34^. Here, we focused on characterizing and quantifying a portion of those cell phenotypes in the brain that may prove to be beneficial for neurodegenerative diseases.

Past chimeric studies report a wide range of methods to quantify donor cell contribution in on and off target tissues, these quantification methods include: fluorescent-activated cell sorting (FACs), qPCR, manual or software-assisted cell counts from tissue histology or a combination of both^24,31–33^. Each method offers advantages and disadvantages. For example, FACs and PCR enable accurate quantification, but require the dissociation of the tissue providing less context about the integration of those cells in space. For quantification methods that use image analysis, there are methodological differences in terms of plane, size, and quality of tissue sections, regions of interest analyzed, the degree of automation, software platforms, cost, and learning curves. These distinct differences indicate that there is an opportunity to explore methods to increase intra- and inter-laboratory reproducibility among chimeric efficiency.

In the current study, we have generated WT intraspecies (mouse-mouse) with microinjection of donor cells into WT blastocysts, for the purpose of studying neural and non-neural cells found within the developing brain. Donor cells formed various neural cell types and contributed to the early developmental brain regions of embryonic day 12.5 mouse-mouse intraspecies chimeras that will become the nucleus basalis, medial septal nucleus, substantia nigra pars compacta, and the dorsal raphe. To explore these brain regions, we have developed a quantification methodology using automated imaging (AI) for two cell types with a tolerable learning curve to collect cell counts of neuronal precursor cells with more consistency. We have also developed an analysis protocol for cell types with simpler morphology (e.g., more round) to collect cell counts for cholinergic/oligodendrocyte precursor and microglia-like cells.

## Methods

### Cell lines: mouse induced pluripotent stem cells

The UMN-3F10 miPSC line was generated to express a transmembrane bound eGFP (Greder et al., 2012). The miPSCs were cultured on irradiated mouse embryonic fibroblasts (#PSC001, R&D Systems, Inc., Minneapolis, MN) in miPSC medium composed of knockout DMEM with 4.5 g/l D-glucose and sodium pyruvate (#10829018, Thermo Fisher Scientific), 10% Gibco^TM^ Knockout^TM^ serum replacement (#A3181501, Thermo Fisher Scientific, Waltham, MA), 10% fetal bovine serum (#SH30071.02HI, HyClone, Logan, UT), 1X Gibco^TM^ MEM non-essential amino acids (#11140050, Thermo Fisher Scientific), 1X Gibco^TM^ GlutaMAX (#35050061, Thermo Fisher Scientific), 0.1 mM 2-mercaptoethanol (#31350010, Thermo Fisher Scientific), 1X Corning^TM^ penicillin/streptomycin (#30002CI, Thermo Fisher Scientific), and 1,000 U/ml ESGRO-LIF (#ESG1107, Millipore Sigma, Burlington, MA). The cells were incubated at 37°C, 5% CO_2_.

### Animals

All research involving mice was approved by the University of Minnesota Institutional Animal Care and Use Committee. Mice had a light cycle of 12:12-h light:dark cycle, with lights on at 7:00 a.m. Surrogate female mice were fed Teklad 2919 diet and all other mice were fed Teklad 2918 diet. Mice were housed in individually ventilated micro-isolator cages on corn-cob bedding and enviro-dri enrichment.

### Mouse zygote isolation

The chimeric embryos were generated, using 3.5-week-old female C57BL6/J mice (Jackson Laboratory, Bar Harbor, ME) that were superovulated using pregnant mare’s serum gonadotropin (5 IU i.p.; National Hormone and Peptide Program, Harbor-UCLA Medical Center, Torrance, CA) at 1:00 p.m. followed 47.5 h later with human chorionic gonadotropin (5 IU i.p.; National Hormone and Peptide Program) and immediately mated with C57BL6/J 4-month-old stud males (Jackson Laboratory, Bar Harbor, ME). Female mice were checked for the presence of a plug the following morning; this is considered embryonic day 0.5 (E0.5). The female mice underwent cervical dislocation, and their ovary, oviduct, and proximal end of the uterine horn were harvested bilaterally and placed in a drop of modified Human Tubal Fluid media (mHTF) (#90126, Irvine Scientific, Santa Clara, CA). Zygotes were extracted from the oviduct of all mice in 3mL of 10mg/mL in hyaluronidase in mHTF (#H4272, Sigma-Aldrich, St. Louis, MO) for no more than 2 min to remove the zygotes from the cumulus-oocyte complex. The zygotes were washed twice in Opti-MEM (#31985062, Thermo Fisher Scientific, Waltham, MA) and then washed twice in mouse human tubal fluid (mHTF) before being transferred to a drop of mHTF covered with mineral oil. Finally, the zygotes were placed in the incubator (37°C, 5% CO2) until they reached the blastocyst stage (E3.5), approximately 72 h later. Around 5 p.m. blastocysts were transferred into the uteri of Avertin anesthetized (225mg/kg) 6-week old CD1 pseudo-pregnant female mice (22-24g; Charles River, Wilmington, MA) two days following the presence of vaginal plug (E2.5) post-mating with 4-month old vasectomized C57BL6/J male mice (Jackson Laboratory, Bar Harbor, ME). The indicated age of the post-implantation embryo was matched to the time-mated CD1 females for all subsequent analyses.

### Preparation of miPSCs for blastocyst injection

miPSC cultures were first washed in 1X PBS without calcium or magnesium (#14190144, Thermo Fisher Scientific), followed by exposing cultures to 0.1 ml/cm^2^ 0.25% Trypsin (#59428C, Sigma Aldrich) for 1 to 3 min at 37°C. Trypsin was neutralized with miPSC media, cell suspension was collected, and centrifuged at 1200 rpms (300g) for 4 min. miPSCs were re-suspended in 1mL of miPSC media, transferred to a 1.5mL microcentrifuge tube, placed on ice for transport to the injection suite, and injected within the next 2 h. Cells were used between passages 10 and 15.

### Mouse blastocyst injection and embryo transfer

The blastocyst injection was performed using eGFP-labeled single cell mouse induced pluripotent stem cells (miPSCs) from the UMN-3F10 line. Cells were suspended with E3.5 mouse blastocysts in a drop of EmbryoMax M2 media (#MR-015-D, Millipore Sigma, Burlington, MA). Approximately 10 miPSCs were microinjected into the blastocoel cavity near the inner cell mass of each blastocyst. After blastocyst complementation, embryos were placed into mHTF drops under mineral oil and incubated at 37°C, 5% CO_2_ for 2-4 h. The complemented embryos were then transferred into the uteri of E2.5 pseudo-pregnant female mice as previously described^22^.

### Mouse embryo extraction for immunohistochemistry

Post-implantation embryos were harvested from pregnant mice that were euthanized via carbon dioxide asphyxiation followed by a pneumothorax between 8 a.m. and 11:30 a.m. on embryonic day 12.5 (E12.5). Uteri were quickly dissected and placed in ice-cold PBS and transported back to the lab on ice for further dissection in a PBS filled petri dish on ice to isolate conceptuses. Embryos were extracted in a fresh petri dish on ice with PBS under a stereomicroscope. Whole embryo images were taken on a fluorescent stereoscope (Leica).

### Immunohistochemistry

Wild type (WT) mouse embryos and complemented mouse-mouse were harvested at E12.5, and immersion-fixed with fresh 4% paraformaldehyde (PFA; #J19943K2, Thermo Fisher Scientific) in PBS overnight at 4°C. Embryos were rinsed in PBS and then cryoprotected by immersion in increasing concentrations of sucrose in PBS up to 30%. The final sucrose immersion was removed from the embryos by dabbing with a paper towel, and samples were embedded in Tissue Plus O.C.T. Compound (#4585, Fisher HealthCare) and snap-frozen on dry ice. Cryosections of 14 μm thickness were cut in the sagittal plane for the brain on a Leica cryostat (Wetzlar, Germany). Slides dried completely prior to being stored at −20°C until the day of staining. Frozen sections were placed on a heating block for 15 to 20 min before antibody staining. Sections were surrounded by a hydrophobic barrier (Vector Labs ImmEDGE hydrophobic barrier pen, Newark, CA). Sections were fixed with 4% PFA for 5 min, followed by three rinses with PBST containing 0.1% Triton X-100 for 5 min. Nonspecific binding was blocked with 10% normal donkey serum (#017-000-121, Jackson ImmunoResearch, West Grove, PA) in PBST. Sections were incubated with primary antibodies overnight at 4°C, then incubated with secondary antibodies and counterstained with DAPI (#D1306, Invitrogen^TM^, Waltham, MA) for 2 h at RT (see antibody information in Table S1&2).

### Imaging

All immunohistological imaging was performed on a Nikon TiE inverted light microscope using a SPECTRA3 light source (Lumencor), a 20X Plan Apo (NA 0.75) objective, and a Hamamatsu ORCA-Flash4.0 V2 camera. The exposure time for a given antibody was selected to maximize the signal-to-noise ratio. Images were acquired and analyzed using the NIS elements advanced research software (Nikon). Images were exported as raw .ND2 files for analysis.

Slides for imaging and immunohistochemistry were selected using the Allen Brain Atlas: Developing Mouse Brain to identify E12.5 brain regions for the regions that will develop into the substantia nigra (SNpc), dorsal raphe nucleus (DRN), nucleus basalis (NB), and the medial septal nucleus (MSN). The Developing Mouse Brain atlas only provides E13.5 atlases, we used this as a guide for determining the correct region for imaging and analysis, based on similar anatomical structures in the E12.5 brains. Due to some variability in slicing of these small samples and the thinness of the sections, we used two sections for most regions as a reference point, which can be found in Figure S1.

### Image Processing: AI-based dopaminergic and serotonergic precursor neuronal cell quantification

Data were sampled from WT (n=5 to 6) and mouse-mouse chimeras (n=5 to 6) depending on the brain region and available sections. Analysis was performed in Nikon NIS-Elements AR software (version 6.10.01) with General Analysis (GA3) and NIS.ai software modules.

For analysis, a 20X stitched image with a z thickness of 14 µm for dopaminergic and cholinergic precursor neurons and 10 µm for microglia and macrophages was captured in the region of interest from all samples. For analysis of the serotonergic precursor neurons, a 20X stitched image with no z thickness was used because of the antibody sensitivity that led to bleaching with a z acquisition. Images with z thickness were then compressed into an EDF for future analysis steps.

For the AI quantification of dopaminergic and serotonergic precursor neurons, images were processed in GA3 with batch processing. Batch processing allows all images to be processed with identical parameters across the samples. For serotonergic neurons, the process was (i) denoising; (ii), local contrast; (iii) rolling ball; and (iv) segment_ai. The SegmentObjects.ai module was trained for 15 iteration on images of WT mice manually segmented with neuronal markers tryptophan hydroxylase 2 (TPH2) for serotonergic precursors to a training loss of 0.0617. This module was then used for automating the detection of individual TPH2 positive cells in the processed images. After the AI-dependent detection process was complete, an operator inspected images for errors and made manual corrections to the detected cells.

Next, in a second batch process with GA3, GFP expression was analyzed without any operator input to eliminate bias. The GFP channel of each analysis was processed with (i) local contrast; (ii) detect regional maxima; and (iii) threshold of 1087 and greater to determine GFP-positive pixels. In another batch process with GA3 for DAPI expression, the DAPI channel was processed with (i) local contrast; and (ii) bright spots, to define DAPI-positive pixels. Each identified neuronal cell was then determined to be GFP-positive or negative based on the overlap with the DAPI-positive signal and set of binary image layers from the AI detection. The number of cells was recorded for each image in Table S3.

A similar process was followed for the dopaminergic precursor neurons, the process was (i) denoising; (ii) local contrast; and (iii) segment_ai. The SegmentObjects.ai module was trained for 5 iteration on images of WT mice manually segmented with neuronal markers tyrosine hydroxylase (TH) for dopaminergic precursors to a training loss of 0.03435. This module was then used for automating the detection of individual TH positive cells in the processed images. After the AI-dependent detection process was complete, an operator inspected images for errors and made manual corrections to the detected cells. The same second batch processing method from the serotonergic analysis was followed for GFP and DAPI channels for the dopaminergic precursors. The number of cells was recorded for each image in Table S4

### Image Processing: Cholinergic precursors and microglia/macrophage quantification using GA3 software

For cholinergic precursor neuron quantification, GA3 module in NIS-Elements, 16-bit multichannel images were processed with batch processing. To process the nucleus basalis region and medial septal nucleus, we followed standard protocol. The nucleic stain, DAPI, was used in the 405 channel, endogenously expressed GFP in the 488 channel, and Olig2 in the 647 channel. Bright spot detection was used in the DAPI channel to identify nuclei. Local contrast, sharpen, and threshold of 500 and greater were used in the 488 channel to identify GFP positive cells. Local contrast, sharpen slightly, and threshold of 375 and greater were used in the 647 channel to identify Olig2 positive cells. These analysis steps were performed automatically and identically for each image. Cell counts that were derived from these image processing steps were counted and tabulated in Table S5.

For microglia and macrophage quantification, GA3 module in NIS-Elements, 16-bit multichannel images were processed with batch processing. The nucleic stain, DAPI, was used in the 405 channel, endogenously expressed GFP in the 488 channel, Lyve1 in the 555 channel, and Iba1 in the 647 channel. Local contrast followed by bright spot detection was used in the DAPI channel to identify nuclei. Local contrast, sharpen, and threshold of 3005 and greater were used in the 488 channel to identify GFP positive cells. Only the threshold of 500 and greater was used in the 555 channel to identify Lyve1 positive cells. Rolling ball, local contrast, threshold of 175 and greater were used in the 647 channel to identify Iba1 positive cells. These analysis steps were performed automatically and identically for each image. Cell counts that were derived from these image processing steps were counted and tabulated in Table S6.

### Statistical analysis

All statistical analysis was performed using GraphPad Prism (GraphPad Software). The cell count data acquired from the image processing section above was compared between the WT and all of a cell type in the mouse-mouse (M-M) animal using an unpaired two-tailed t-test.

Further analysis was conducted with a repeated measures one-way ANOVA and multiple comparison’s test, Tukey test, comparing the M-M chimeras all of a cell type, GFP only of that cell type, and host only of that cell type. Donor and host cell contribution was analyzed with a paired two-tailed t-test. Some analyses had values that appeared to be outliers; these data were kept due to the diversity of chimeric contribution that can be seen when generating WT chimeras. Results were reported as mean ± the standard error of the mean (SEM). Differences were regarded as statistically significant where the p-value was <0.05.

## Results

### Donor GFP-labeled stem cells generate mouse-mouse chimeric embryos with chimeric brains

Mouse-mouse chimeras were generated by performing superovulation and timed-mating to extract zygotes. Next, EGFP-labeled miPSCs were injected into wild type (WT) blastocysts at embryonic day (E) 3.5 and transferred into surrogate pseudopregnant dams (Figure 1a). Embryos collected at E12.5 generally showed robust GFP expression throughout the body (Figure 1b).

**Figure 1.**
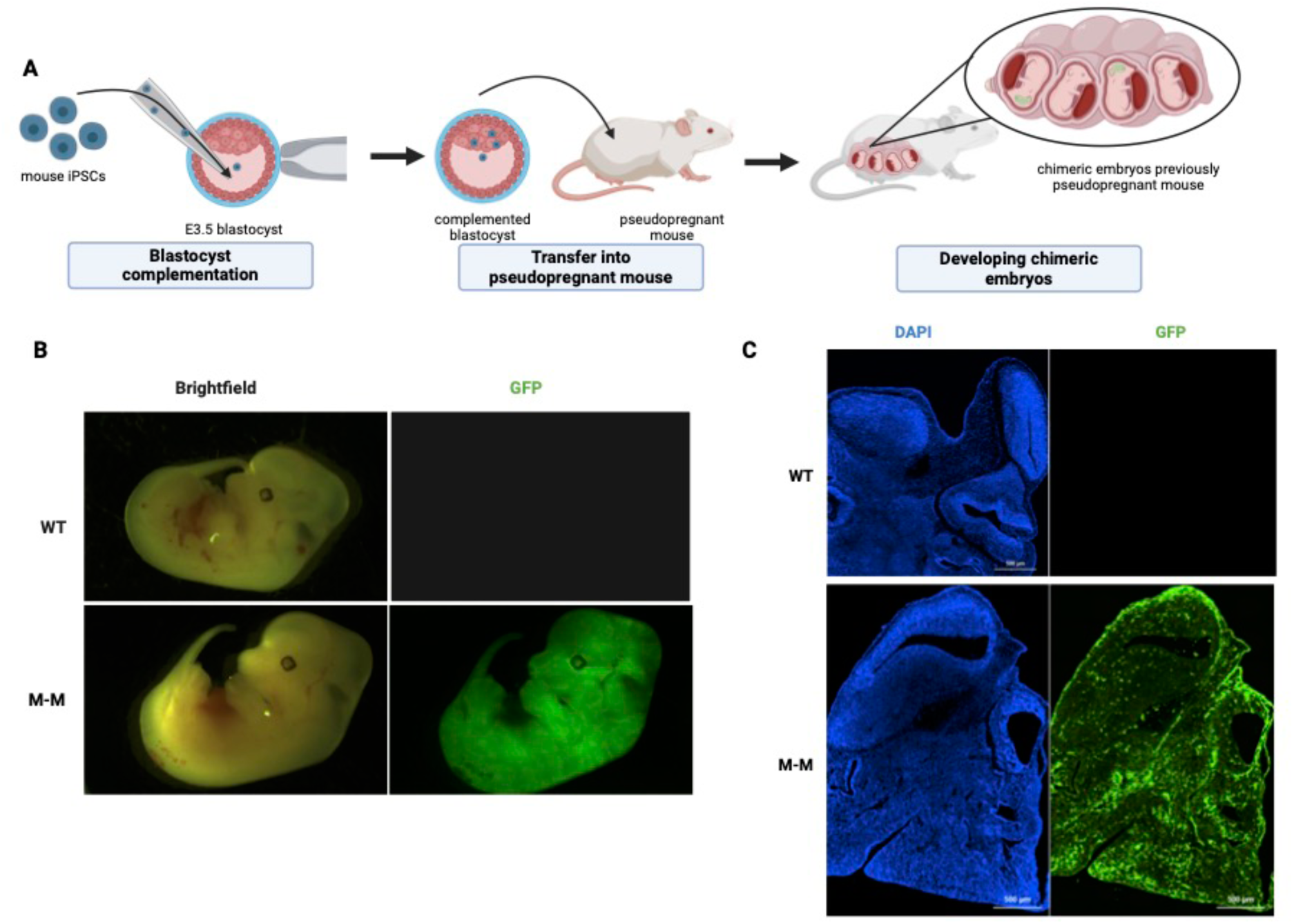
GFP-labeled donor cell robust expression and contribution in a WT M-M chimera (a) Overview schematic of generating WT M-M chimeras (b) Representative images of E12.5 WT and M-M whole embryos with robust GFP expression in the M-M chimera. (c) Representative images of 14 µm sections of E12.5 WT and M-M heads to show the presence of GFP-labeled donor cells throughout the M-M head and brain regions.

**Figure 2.**
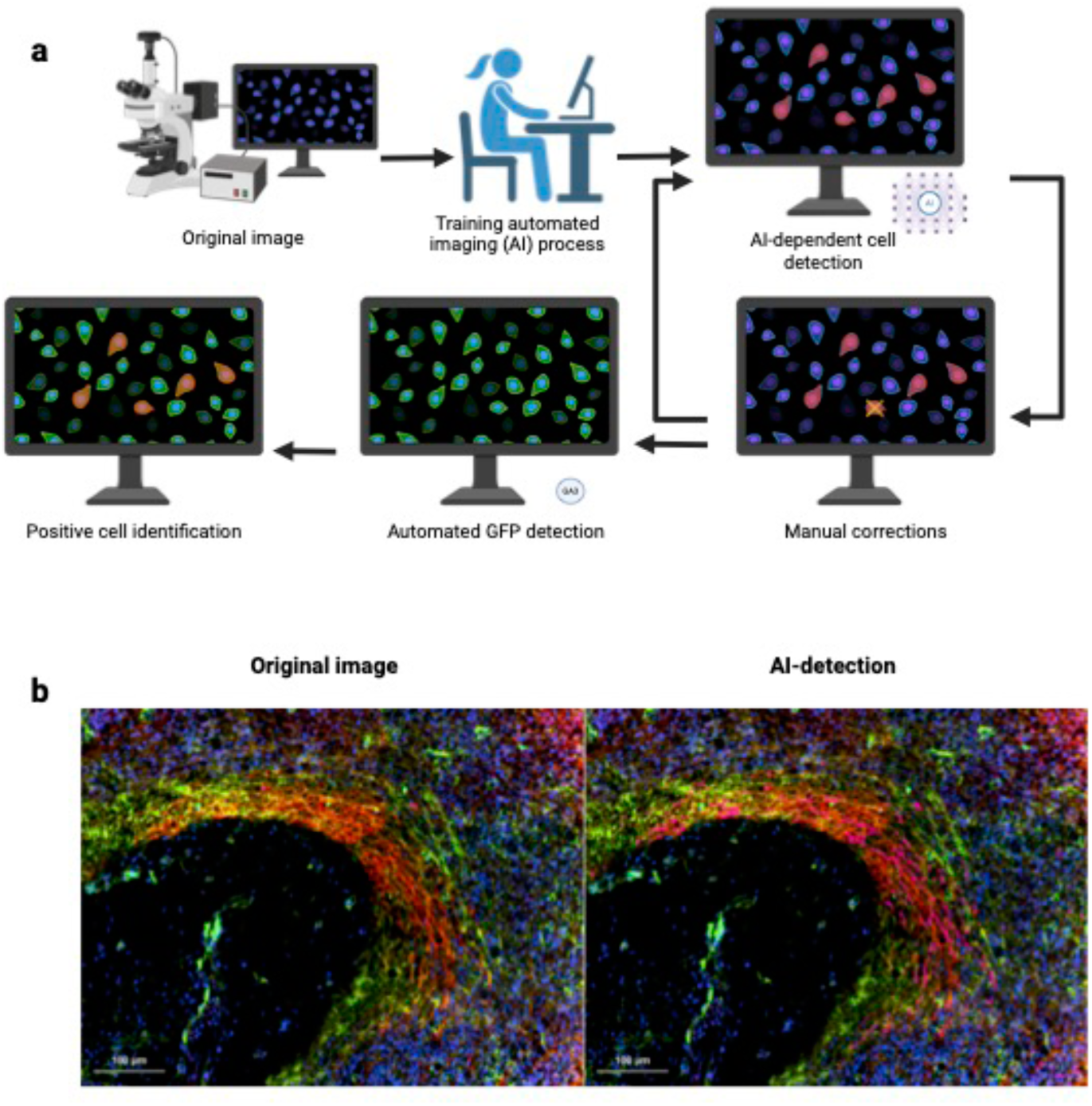
Automated cell detection method. (a) Overview of automated neuronal or microglia donor cell analysis approach. Steps 1-3 and 5-6 were performed without an operator, so they were identical and automatic after initial setup. Initial Step 2 setup was completed by an operator to train AI, after setup this was completed without an operator. Step 4 was performed by an operator. (b) Representative images of the original image on the left contains GFP (green) from donor cells and TH (red) for the dopaminergic precursor cells. The AI-dependent detection contains the GFP (green) from the donor cells, and now the dopaminergic precursor cells (red) are outlined in pink. Scale bar, 100 µM.

Upon further analysis, donor-derived GFP cells were observed in the mouse-mouse chimera brain compared to a WT brain with no GFP donor cells (Figure 1c). The GFP-labeled cells contributed to both early neuronal and non-neuronal cells in the mouse-mouse chimeras with robust donor cell contribution (Figure 1c). Previous studies demonstrated donor cell contribution to the brain via blastocyst complementation with mouse stem cells^30,31^. Our results confirmed the previous findings, and demonstrated on a broad scope that many cell types within the brain can be derived from the donor cells.

### Donor stem cells contribute to relevant developing chimeric mouse-mouse brain regions

Donor cell contribution appeared throughout the mouse-mouse (M-M) brain (Figure 1b & 1c), in support of previously reported studies^31^. In the developing M-M chimeras, GFP-labeled cells were found in the developing brain regions of the dorsal raphe nucleus (DRN), substantia nigra pars compacta (SNpc), nucleus basalis (NB), and medial septal nucleus (MSN) (Figure 3–6). Under further analysis, we have begun to characterize cell types in these developing brain regions that may be important for cellular replacement or transplantation therapies for neurodegenerative diseases.

**Figure 3.**
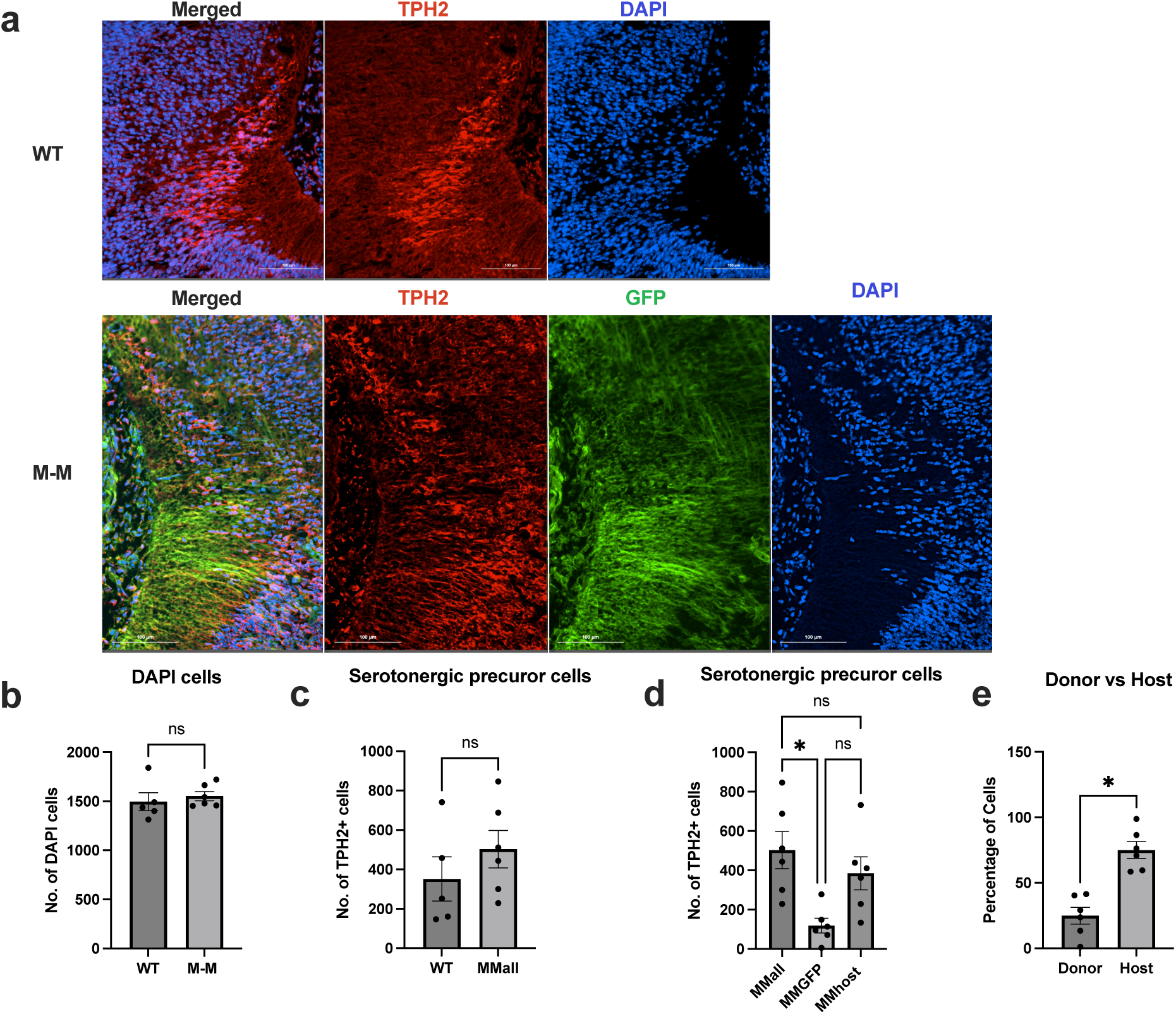
Quantification of serotonergic precursor cells in the E12.5 chimeric M-M brain. (a) Representative images from E12.5 WT and M-M developing region of the DRN, stained with TPH2–a serotonergic neuronal marker. (b) Total number of DAPI cells detected in the brain regions analyzed (unpaired t-test, p=0.5839). (c) Total number of TPH2-positive cells detected in the WT (n=5) and M-M (n=6) chimeras in the analyzed region (unpaired t-test, p=0.3268). (d) Comparison of the total number of TH positive cells detected within each M-M (n=6) chimera that were also GFP positive (MMGFP) and GFP negative (MMhost), with fewer serotonergic cells detected that were GFP positive (repeated measures one-way ANOVA, p=0.0065). (e) Percentage of donor and host cell contribution to brain region under analysis with fewer donor-derived cells contributing to serotonergic cells found in the developing DRN (n=6, paired t-test, p=0.0112).

### AI quantification of serotonergic and dopaminergic neuronal precursors derived from GFP- labeled donor cells

We adapted an AI quantification workflow to detect the level of co-labeled neuronal precursor cells^35^ (Figure 2a). The AI-dependent cell detection allowed for the detection of a complex cell morphology with a more consistent and reliable cell counting system while avoiding subjective human operator errors (Figure 2b). Immunological fluorescent identification of serotonergic and dopaminergic neuronal precursors was performed with tryptophan hydroxylase 2 (TPH2) and tyrosine hydroxylase (TH), respectively.

The developing DRN region showed that WT and M-M chimeras had approximately the same quantity of total cells and total serotonergic precursors detected in the analyzed developing brain regions (Figure 3a-c). The serotonergic precursor cells were derived from donor stem cells, observed by the co-localization of the GFP-labeled donor cells and TPH2, a serotonergic neuron marker. However, the donor derived cell count mean of 118.7 was lower than that of the host derived cell count mean of 384.3 (Figure 3a & 3d). When examining the proportion of donor-derived compared to host-derived TPH2-positive cells, there was a significantly lower percentage of donor-derived TPH2-positive cells (Figure 3d & 3e). These observations suggested that exogenic serotonergic precursors derived from GFP-labelled donor cells can be generated in an intraspecies chimera.

Further examination of the SNpc, using TH to identify dopaminergic precursors, showed that WT and M-M chimeras had approximately the same quantity of total cells and total dopaminergic precursors detected in the analyzed developing brain regions (Figure 4a-c). The dopaminergic precursor cells were derived from donor stem cells, as evident by the co-localization of the GFP-labeled donor cells and TH (Figure 4a & 4d). We also observed dopaminergic precursors in the M-M chimeras that originated from host cells (Figure 4d). The quantity of donor-derived TH-positive cells was about half of the total dopaminergic precursors detected (Figure 4d & 4e). The degree of donor and host cell contribution of TH-positive cells showed no difference between the two groups, with ∼ 50% coming from the donor and host cells (Figure 4e). Together, these observations suggested that both serotonergic and dopaminergic precursors were derived from GFP-labeled donor cells. However, dopaminergic precursors have more equal proportions of cells derived from donor and host cells compared to serotonergic, as quantified with the use of AI-dependent cell detection methods. This suggests the percentage of donor cell contribution may be dependent on a given brain region and/or cell type.

**Figure 4.**
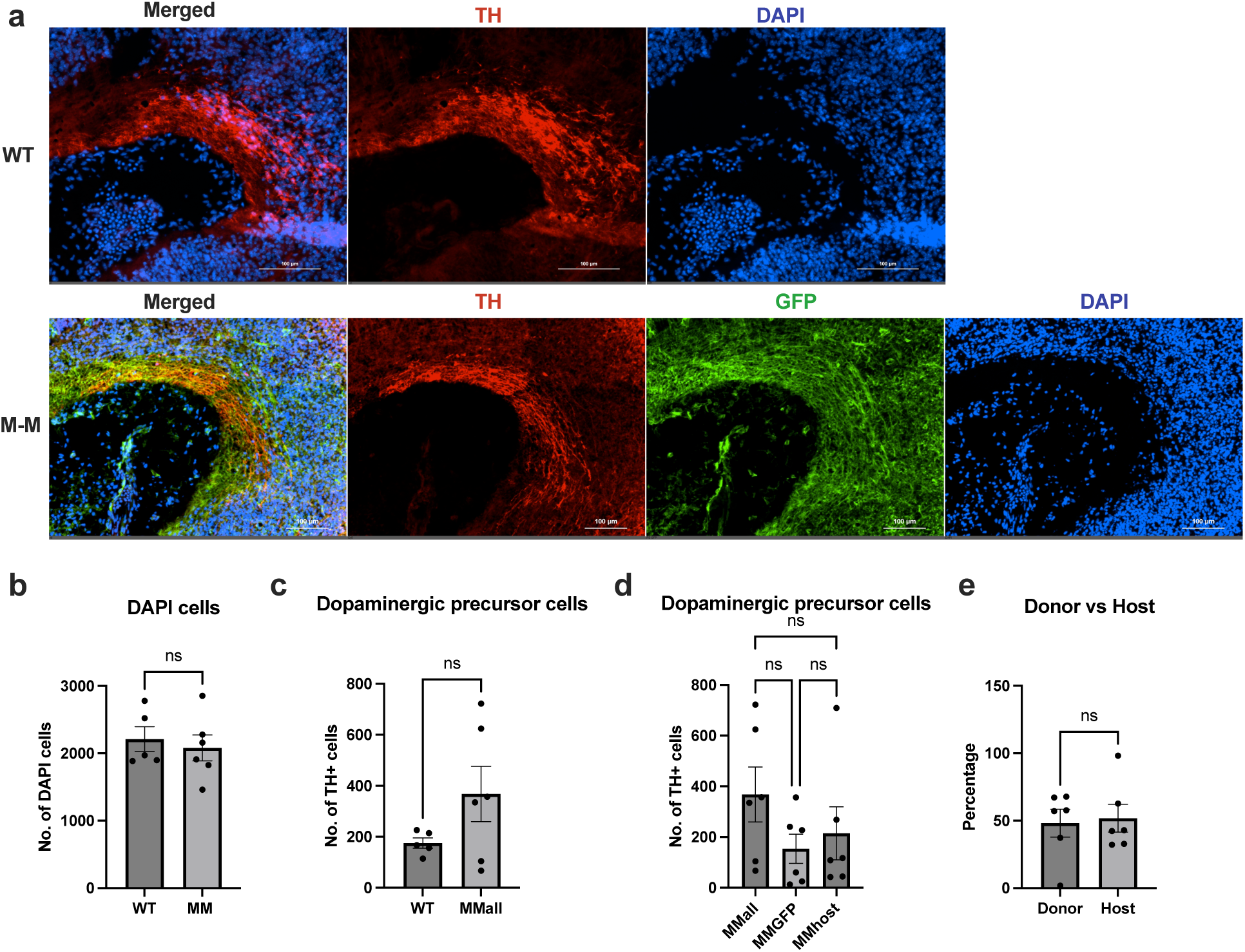
Quantification of dopaminergic precursor cells in the E12.5 chimeric M-M brain. (a) Representative images from E12.5 WT and M-M developing region of the SNpc, stained with TH–a dopaminergic neuronal marker. (b) Total number of DAPI cells detected in the brain regions analyzed (unpaired t-test, p=0.6436). (c) Total number of TH-positive cells detected in the WT (n=5) and M-M (n=6) chimeras in the analyzed region (unpaired t-test, p=0.1462). (d) Comparison of the total number of TH positive cells detected within each M-M (n=6) chimera that were also GFP positive (MMGFP) and GFP negative (MMhost) (repeated measures one-way ANOVA, p=0.1718). (e) Percentage of donor and host cell contribution to brain region under analysis (n=6, paired t-test, p=0.8639).

### Quantification of cholinergic neuronal precursors derived from GFP-labeled donor cells

To examine the developing NB and MSN brain regions in the WT and M-M chimeras, we analyzed brain regions that had approximately the same quantity of total cells and total cholinergic precursors for the given brain region (Figures 5a-c & 6a-c). An early marker for cholinergic cell development, Olig2, was selected to identify possible cholinergic neurons expressed at E12.5^36^. The NB contained cholinergic precursor cells derived from donor stem cells, as observed by the co-localization of the GFP-labeled donor cells and Olig2 (Figure 5a & 5d). Cholinergic precursors in the M-M NB also originated from host cells (Figure 5d). We noted a greater cell count of donor-derived Olig2-positive cells than that of the Olig2-positive host cells (Figure 5d & 5e). These donor and host cell counts are almost significantly different from the total number of Olig2-positive cells that should develop within this region in the M-M chimera (Figure 5d).

**Figure 5.**
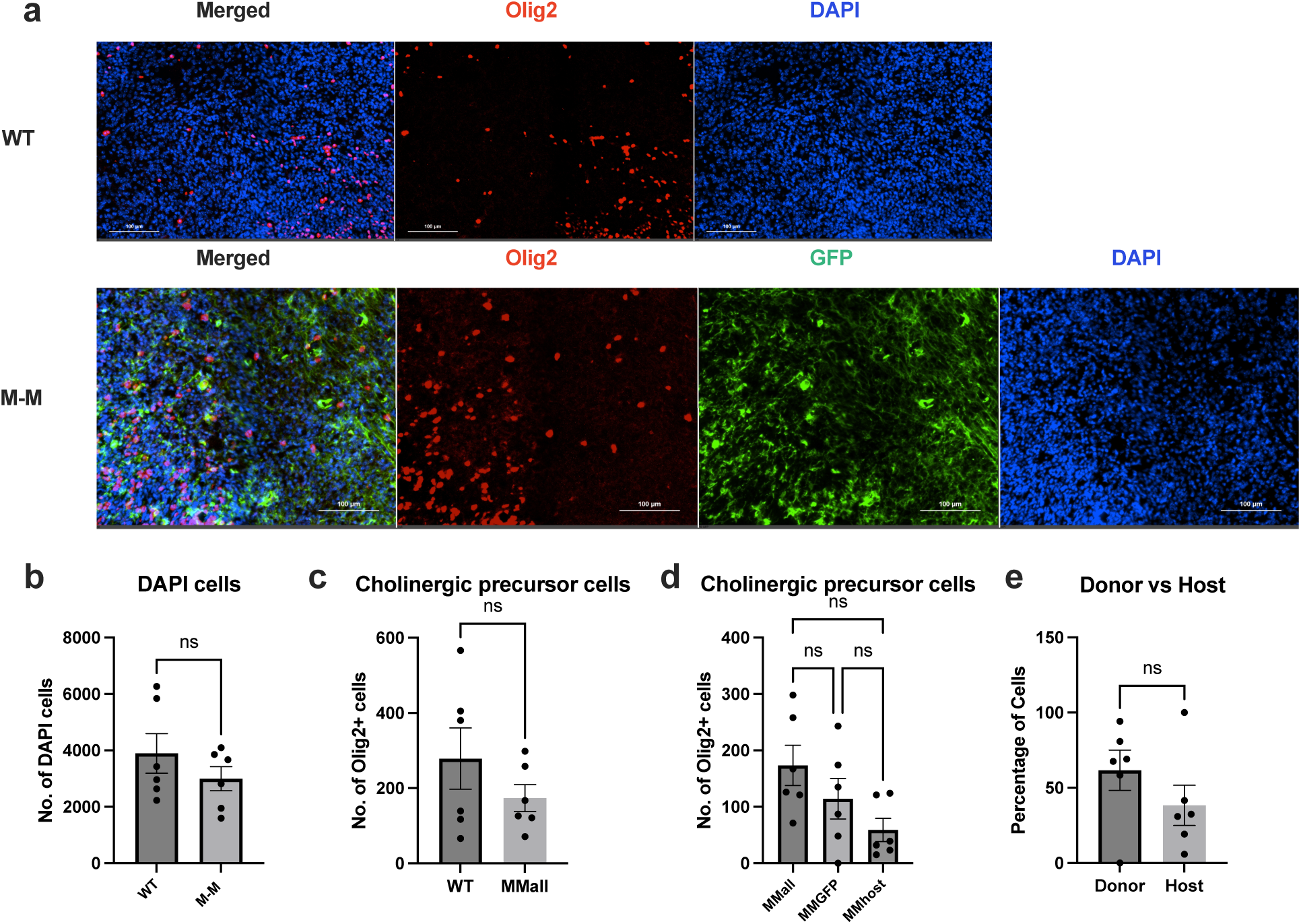
Quantification of cholinergic precursor cells in the E12.5 chimeric M-M developing NB brain region. (a) Representative images from E12.5 WT and M-M developing region of the NB, stained with Olig2–a cholinergic precursor marker. (b) Total number of DAPI cells detected in the brain regions analyzed (unpaired t-test, p=0.3016). (c) Total number of Olig2-positive cells detected in the WT (n=6) and M-M (n=6) chimeras in the analyzed region (unpaired t-test, p=0.2642). (d) Comparison of the total number of Olig2-positive cells detected within each M-M (n=6) chimera that were also GFP positive (MMGFP) and GFP negative (MMhost) (repeated measures one-way ANOVA, p=0.0583). The donor derived cells and host cells are nearly different from the total number of Olig2-positive cells that should develop in that space (Tukey’s comparisons test: MMall vs MMGFP, p=0.0737; MMall vs MMhost p=0.0538). (e) Percentage of donor and host cell contribution to brain region under analysis (n=6, paired t-test, p=0.4210).

The MSN also contained cholinergic precursor cells derived from donor stem cells, as seen with the co-localization of the GFP-labeled donor cells and Olig2 (Figure 6a & 6d). Cholinergic precursors in the M-M MSN also originated from host cells (Figure 6d). The quantity of donor-derived Olig2-positive cells was lower than that of the Olig2-positive host cells, with no significant difference between the cell counts of the donor and host cells (Figure 6d & 6e). A comparison of the donor-derived cells, host-derived cells, and total number of counted cells showed that the Olig2-positive donor cell count mean and Olig2-positive host cell count mean were significantly different than the total number of Olig2-positive cells present in the M-M chimera (Figure 6d). The degree of donor and host cell contribution to Olig2-positive cells showed no difference between the two groups, with a slight trend in favor of more cells contributed from the host (Figure 6e). Together these observations suggested that cholinergic precursors in the developing NB and MSN can be derived from GFP-labeled donor cells, but the number of GFP-derived cells detected within these regions may vary depending on the developing brain region.

**Figure 6.**
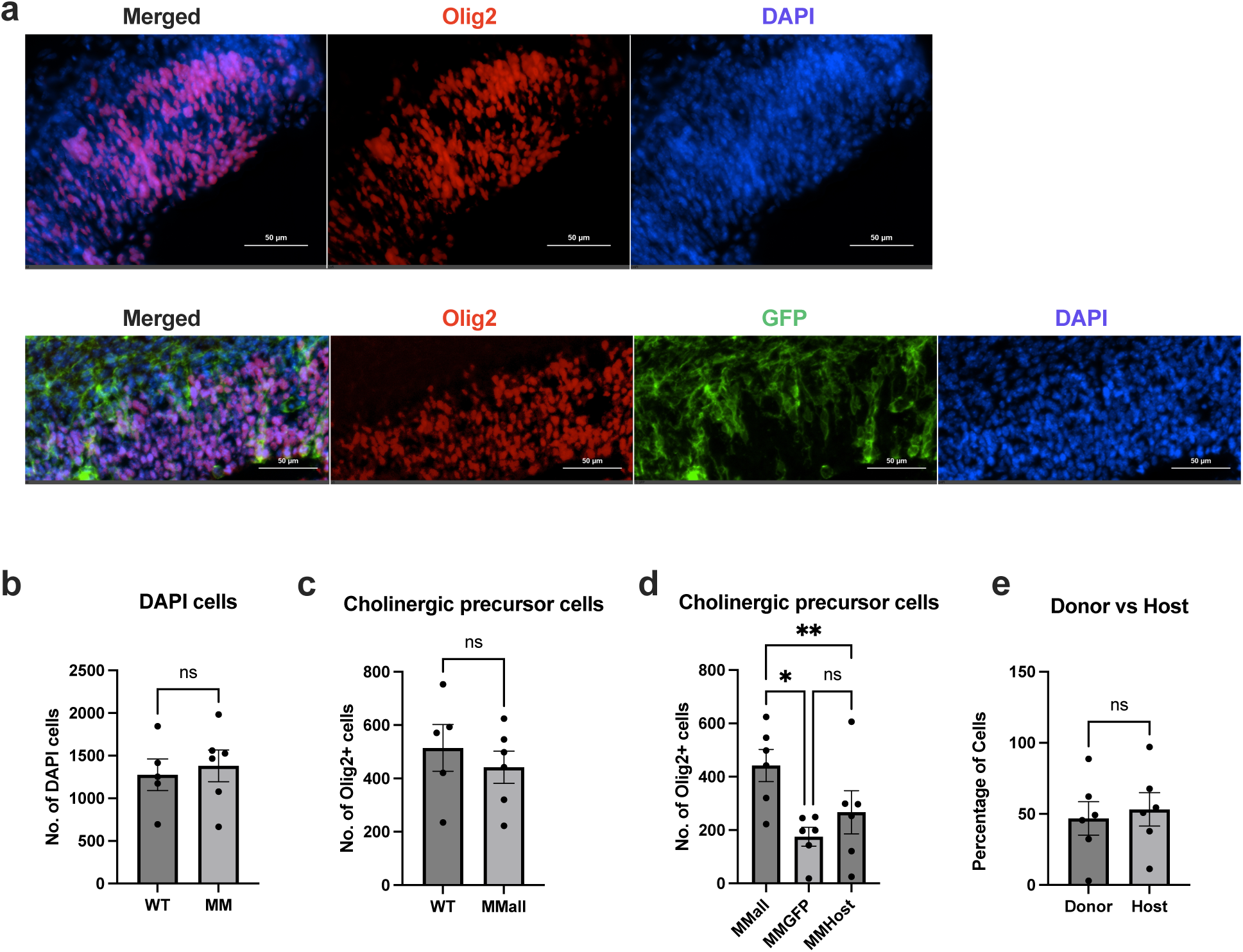
Quantification of cholinergic precursor cells in the E12.5 chimeric M-M developing MSN brain region. (a) Representative images from E12.5 WT and M-M developing region of the MSN, stained with Olig2–a cholinergic precursor marker. (b) Total number of DAPI cells detected in the brain regions analyzed (unpaired t-test, p=0.7033). (c) Total number of Olig2-positive cells detected in the WT (n=5) and M-M (n=6) chimeras in the analyzed region (unpaired t-test, p=0.5012). (d) Comparison of the total number of Olig2-positive cells detected within each M-M (n=6) chimera that were also GFP positive (MMGFP) and GFP negative (MMhost) (repeated measures one-way ANOVA, p=0.0603). The donor derived cells and host cells were significantly different from the total number of Olig2-positive cells that should develop in that space (Tukey’s comparisons test: MMall vs MMGFP, p=0.0479; MMall vs MMhost p=0.0096). (e) Percentage of donor and host cell contribution to brain region under analysis (n=6, paired t-test, p=0.7979).

### Quantification of microglia-like cells and macrophage-like cells derived from GFP-labeled donor cells

As previously noted, we observed robust expression of GFP-labeled donor cells in neuronal and non-neuronal cells (Figure 1c). Using a similar brain region across all samples for immunofluorescence and analysis, we generated microglia-like cells derived from GFP-labeled donor cells (Figure 7a & 7c), as observed by the co-localization of GFP and IBA1, microglia and macrophage cell surface marker^37^. To distinguish microglia-like cells from macrophage-like cells, we stained samples with LYVE1, an endothelial and perivascular macrophage marker^38,39^. Therefore, cells double-positive for IBA1 and LYVE1 were macrophages, and cells positive for IBA1 and not LYVE1 were microglia-like cells. A similar quantity of microglia-like cells and macrophage-like cells were detected in the WT and M-M chimeras (Figure 7b & 7e). Both donor and host-derived microglia-like cells and macrophages were observed, with greater numbers of cells derived from the GFP-labeled donor cells (Figure 7c & 7f). There was a greater degree of donor cell contribution to microglia-like cells and macrophages compared to host cell contribution (Figure 7d & 7g). Notably, the donor-derived macrophage-like cells were significantly more abundant compared to the host macrophage-like cells (Figure 7g). The microglia-like cells trended in this direction but had one animal that had much fewer donor-derived microglia compared to host cells. These findings suggested GFP-labeled donor cells injected into an early-stage blastocyst, more readily contribute to microglia-like cells and macrophages in intraspecies chimeras, with one outlier with little donor cell contribution.

**Figure 7.**
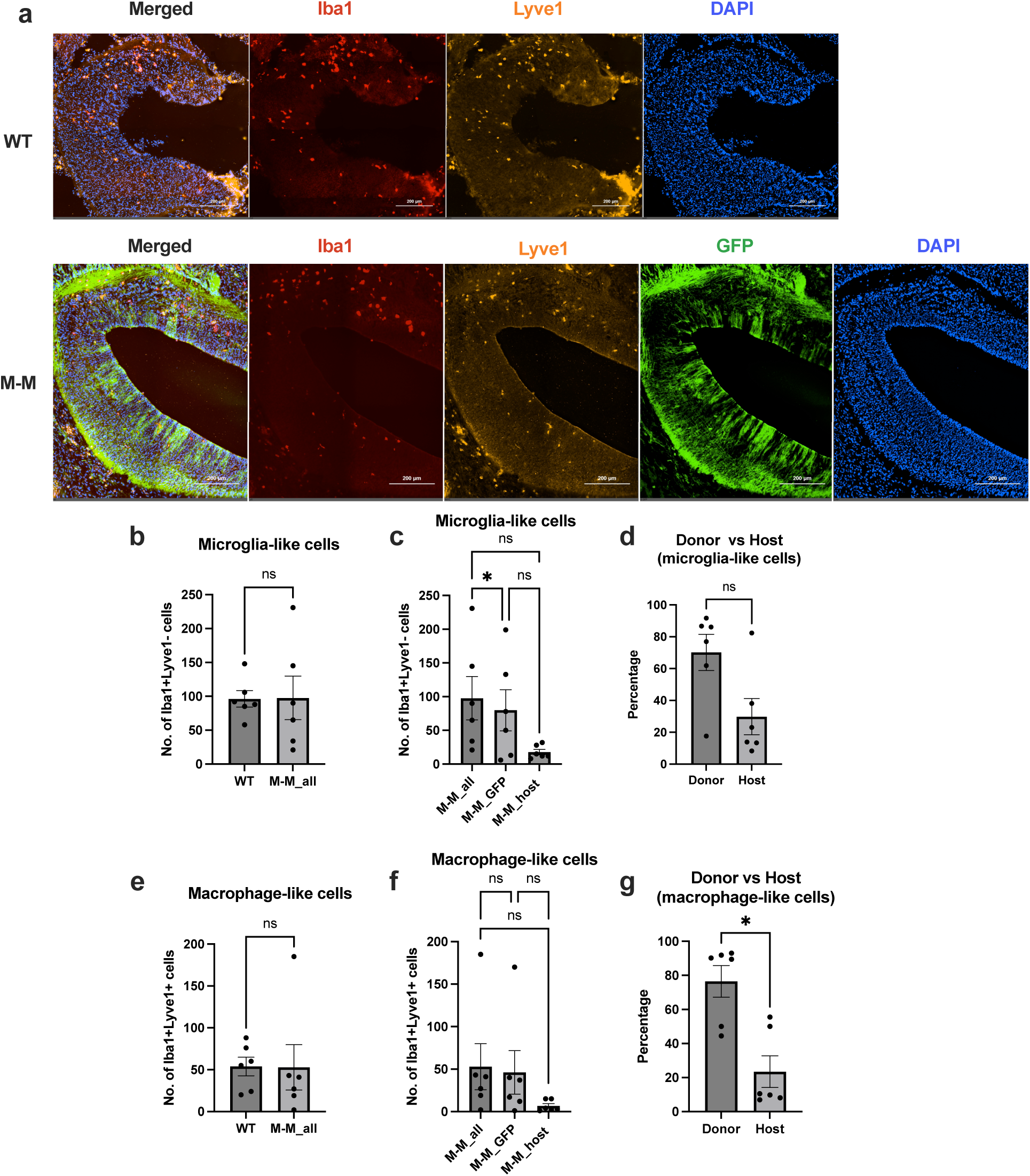
Quantification of microglia-like cells and macrophage-like cells. (a) Representative images of WT and M-M chimera, showing staining of Iba1 (red), Lyve1 (orange), GFP (green), and DAPI (blue). (b) Number of Iba1+Lyve1-cells detected in the WT (n=6) and M-M (n=6) chimeric animals (unpaired t-test, p=0.9661). (c) The total number of microglia-like cells detected (MM_all) cells detected in the M-M chimera (n=6), along with the number of donor cells that expressed Iba1+Lyve1-GFP+ (MM_GFP) and the number of host cells that expressed Iba1+Lyve1-(MM_host) (repeated measures one-way ANOVA, p=0.0584). (d) The percentage of microglia-like cells derived from the donor GFP-labeled cells compared to the host cells (n=6, paired t-test, p=0.1357). (e) Number of Iba1+Lyve1+ cells detected in the WT (n=6) and M-M (n=6) chimeric animals (unpaired t-test, p=0.9735). (f) The total number of macrophage-like cells detected (MM_all) cells detected in the M-M chimera (n=6), along with the number of donor cells that expressed Iba1+Lyve1+GFP+ (MM_GFP) and the number of host cells that expressed Iba1+Lyve1-(MM_host) (repeated measures one-way ANOVA, p=0.1419). The GFP-derived donor cell count showed that it was different from the control cell count of microglia-like cells in M-M chimera (Tukey’s comparisons test: MMall vs MMGFP, p=0.0150). (g) The percentage of macrophage-like cells derived from the donor GFP-labeled cells compared to the host cells (n=6, paired t-test, p=0.0358).

## Discussion

In this study, we have shown that characterization of intraspecies chimeras varies depending on cell type and brain region to produce chimeric M-M brains. Using donor and host from the same species, typically encourages higher levels of chimerism when using blastocyst complementation^24^. We found that the average percentage of donor cell contribution ranged from 50-80% depending on the cell type at E12.5 with microglia-like cells and macrophages having higher donor cell contribution compared to the neuronal precursors. The neuronal precursor contributions aligned with adult cortex and midbrain contribution as previously observed^31^.

Despite the same species for the donor cells and host embryo, there was still large variability in donor-cell contribution in some cases within a cell type. For example, one M-M chimera had ∼17% of microglia-like cells derived from donor cells and 83% derived from host cells. Another M-M chimera had ∼ 2% dopaminergic precursor cells derived from donor cells and 98% derived from host cells. Thus, there appear to be some hurdles that exist within the blastocyst complementation field even for intraspecies chimeras, including the brain.

Our results underscore the significance of having a baseline for neuronal and non-neuronal cell types that can be produced in the developing intraspecies M-M brain. We found that GFP-labeled donor cells contributed less to overall levels of chimerism in neuronal precursors and specific developing brain regions than in non-neuronal cells, like the early macrophage-like cells. This suggests that there may be barriers that influence the success of a donor cell integrating into a particular cell type in a given region. For example, microglia and macrophage cells arise from the mesoderm germ layer that originates from the yolk sac, while neurons and other glial cells arise from the ectoderm^40,41^. Thus, how and where these donor cells integrate during gastrulation influences part of their success in developing further along in the host. Additionally, there may be more conserved developmental cues in the generation of these myeloid lineages than in neurons.

More recent studies showed that WT chimeras can generate functional cells^32,42^. In some cases, these donor-derived cells have been shown to treat certain diseases such as liver fibrosis, and restore normal hepatocyte morphology and function^42^. This suggests that WT donor-derived may be sufficient to treat degenerative diseases, especially diseases of the brain. A more recent study focused on contribution of donor cells to various brain regions in WT rat-mouse chimeric brain^32^, but did not focus on intraspecies brain development, early embryo brain development, and developing cell types within the brain. Our study aims to generate a foundation to characterize some of these developing neural cells, as a possible future means of cellular therapy.

Cell transplantation therapies have successfully engrafted and reversed cognitive and behavioral deficits of neurodegenerative disease models^8,18,43^. Cholinergic neurons extracted from the E14-E15 medial septal nucleus of rats have shown to engraft, express acetyl cholinesterase, and improve memory^44,45^. More recent studies have aimed to use stem-cell derived cholinergic neurons, which have also been shown to survive post transplantation, express choline acetyltransferase, and form functional synapses^46,47^. Fetal rat serotonergic neurons have also survived post-transplantation, innervated, and may restore serotonin levels^48–50^. Dopamine precursor cells from human fetal brain and iPSC-derived cells were shown to engraft and ameliorate movement disorders in PD patients^11–16^. As for non-neuronal cells, like microglia, that are implicated in neurodegenerative disease pathologies, these cells may ameliorate pathology and restore neurological function^43,51^. Our study identifies some of these neuronal precursors such as dopaminergic, serotonergic, and cholinergic cells, and non-neuronal cells like microglia-like cells in the WT mouse-mouse chimeric brain. The use of the characterized cells may be beneficial in future cell transplantation studies.

In summary, we have characterized plausible neuronal and non-neuronal precursor cells generated through blastocyst complementation that could be used in future cell transplantation therapies for neurodegenerative diseases. Our intraspecies complementation strategy focused on the first steps of characterizing these cell types and proportions of donor-cell contribution in these developing brain regions. A limitation of this study is that it does not address the more complex goal of interspecies chimeras; however, a more simplified view of intraspecies chimeras provides a baseline for future studies.

## Acknowledgements

The authors thank the University of Minnesota Imaging Core for imaging equipment used for this research; Alicia Strtak from Nikon Instruments for additional software training; Figures 1 & 2 were created with the help of BioRender.com

## Data Availability Statement

The raw data that support the findings of this study are available from the corresponding author upon reasonable request.

