## Supplementary material for "Characterization of the intraspecies chimeric mouse brain at embryonic day 12.5": Figure S1

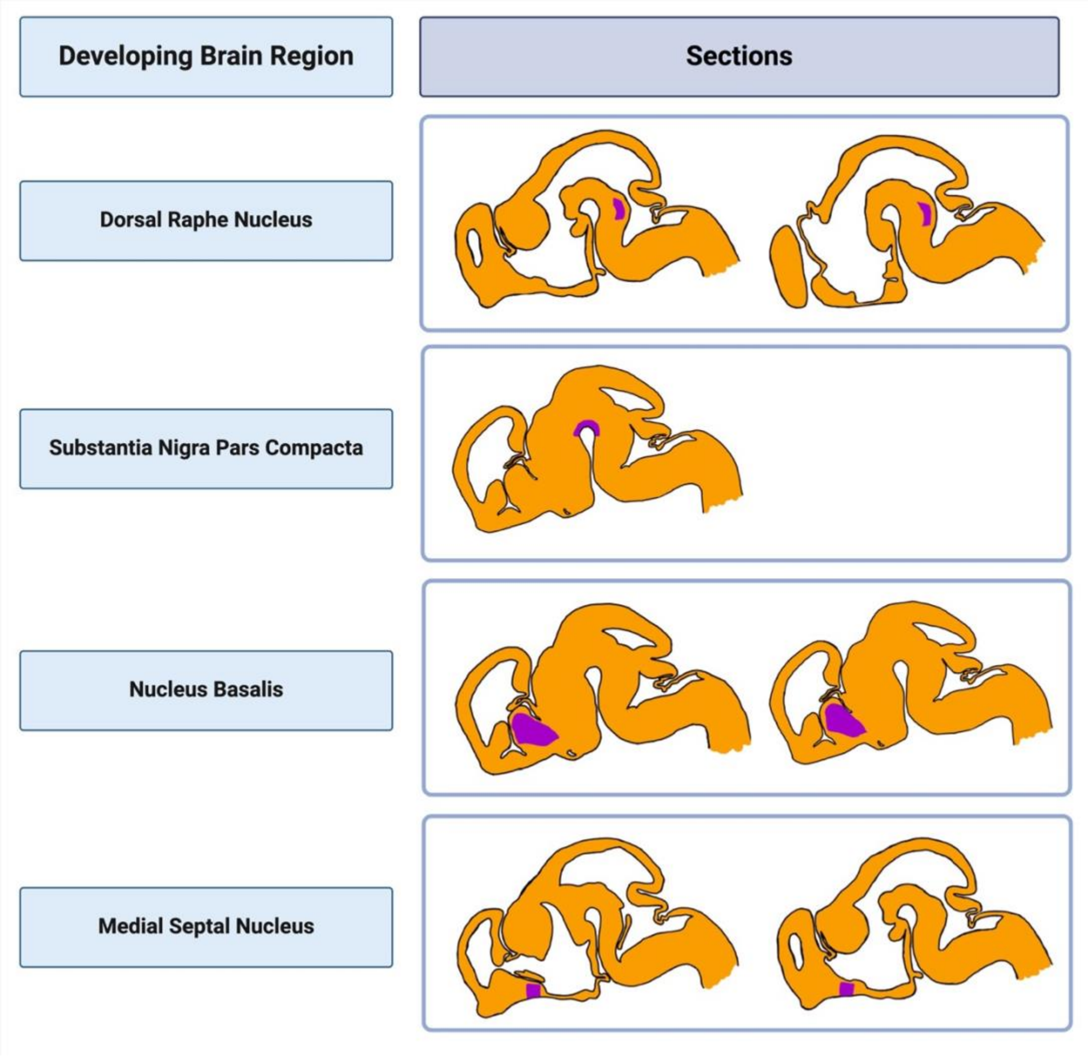


**Figure S1.** Slide selection and brain region for analysis in E12.5 brains based on the Allen Brain Atlas: Developing Mouse Brain. For the adult brain regions (listed on the left), their corresponding developing region was located in the Developing Mouse Brain atlas on one or two sections, and used for future imaging and analysis. These developing areas are noted in purple on the sections. The DRN developing region is the periventricular stratum of basal medial part of rhombomere 1 (r1BM). The SNpc developing region includes the mantle zone of basal plate of mesomere 1 (m1B) and the mantle zone of basal plate of prosomere 1 (p1B), 2 (p2B), and 3 (p3B). The NB developing region is part of the central subpallium. The MSN developing region comes from the mantle zone of diagonal part of septum.
