## Supplementary material for "Characterization of the intraspecies chimeric mouse brain at embryonic day 12.5": Table S1

| Antibody | Species | Dilution | Company (catalog #) |
|---|---|---|---|
| Olig2 | Rabbit | 1:200 | Abcam (ab109186) |
| TPH2 | Rabbit | 1:500 | Invitrogen (PA1-778) |
| TH | Rabbit | 1:500 | Chemicon (AB152) |
| IBA1 | Rabbit | 1:1000 | Fujifilm Wako (019-19741) |
| LYVE-1 | Goat | 1:200 | R&D systems (AF2125) |
