## Supplementary material for "Characterization of the intraspecies chimeric mouse brain at embryonic day 12.5": Table S2

| Antibody | Species | Dilution | Company (catalog #) |
|---|---|---|---|
| Donkey Anti-Goat IgG H&L (Alex Fluor® 555) | Goat | 1:1000 | Abcam (ab150130) |
| Donkey F(ab')2 Anti-Rabbit IgG H&L (Alexa Fluor® 647) preadsorbed | Rabbit | 1:1000 | Abcam (ab181347) |
