## Supplementary material for "Characterization of the intraspecies chimeric mouse brain at embryonic day 12.5": Table S3

| Sample Type | DAPI | TPH2 all | TPH2 MM | TPH2 Host |
|---|---|---|---|---|
| WT | 1491 | 160 | n/a | n/a |
| WT | 1841 | 252 | n/a | n/a |
| WT | 1438 | 147 | n/a | n/a |
| WT | 1401 | 741 | n/a | n/a |
| WT | 1314 | 458 | n/a | n/a |
| MM | 1664 | 688 | 278 | 410 |
| MM | 1457 | 846 | 114 | 732 |
| MM | 1477 | 511 | 149 | 362 |
| MM | 1452 | 444 | 5 | 439 |
| MM | 1720 | 300 | 71 | 229 |
| MM | 1542 | 229 | 95 | 134 |
