## Supplementary material for "Characterization of the intraspecies chimeric mouse brain at embryonic day 12.5": Table S4

| Sample Type | DAPI | TH all | TH MM | TH Host |
|---|---|---|---|---|
| WT | 1886 | 114 | n/a | n/a |
| WT | 1899 | 226 | n/a | n/a |
| WT | 2522 | 167 | n/a | n/a |
| WT | 1970 | 214 | n/a | n/a |
| WT | 2778 | 156 | n/a | n/a |
| MM | 1460 | 357 | 240 | 117 |
| MM | 1910 | 104 | 60 | 44 |
| MM | 2158 | 624 | 356 | 268 |
| MM | 2279 | 722 | 13 | 709 |
| MM | 2854 | 335 | 227 | 108 |
| MM | 1827 | 67 | 25 | 42 |
