## Supplementary material for "Characterization of the intraspecies chimeric mouse brain at embryonic day 12.5": Table S5

| Cholinergic region | Sample Type | DAPI | Olig2 all | Olig2 MM | Olig2 Host |
|---|---|---|---|---|---|
| NB | WT | 3427 | 566 | n/a | n/a |
| NB | WT | 2634 | 380 | n/a | n/a |
| NB | WT | 2964 | 117 | n/a | n/a |
| NB | WT | 5846 | 66 | n/a | n/a |
| NB | WT | 2232 | 139 | n/a | n/a |
| NB | WT | 6269 | 405 | n/a | n/a |
| NB | MM | 1597 | 258 | 243 | 15 |
| NB | MM | 1948 | 167 | 135 | 32 |
| NB | MM | 4094 | 298 | 174 | 124 |
| NB | MM | 3859 | 126 | 87 | 39 |
| NB | MM | 2817 | 121 | 0 | 121 |
| NB | MM | 3671 | 71 | 48 | 23 |
| MSN | WT | 1844 | 253 | n/a | n/a |
| MSN | WT | 1416 | 753 | n/a | n/a |
| MSN | WT | 1173 | 421 | n/a | n/a |
| MSN | WT | 1253 | 593 | n/a | n/a |
| MSN | WT | 694 | 571 | n/a | n/a |
| MSN | MM | 1465 | 498 | 245 | 253 |
| MSN | MM | 1079 | 222 | 197 | 25 |
| MSN | MM | 1527 | 442 | 143 | 299 |
| MSN | MM | 1560 | 546 | 249 | 297 |
| MSN | MM | 1983 | 624 | 18 | 606 |
| MSN | MM | 666 | 320 | 199 | 121 |
