## Supplementary material for "Characterization of the intraspecies chimeric mouse brain at embryonic day 12.5": Table S6

| Sample Type | DAPI | IBA1+ LYVE1- all | IBA1+ LYVE1- MM | IBA1+ LYVE1- Host | IBA1+ LYVE1+ all | IBA1+ LYVE1+ MM | IBA1+ LYVE1+ Host |
|---|---|---|---|---|---|---|---|
| WT | 3136 | 88 | n/a | n/a | 24 | n/a | n/a |
| WT | 6428 | 85 | n/a | n/a | 60 | n/a | n/a |
| WT | 5410 | 148 | n/a | n/a | 88 | n/a | n/a |
| WT | 6648 | 58 | n/a | n/a | 55 | n/a | n/a |
| WT | 4436 | 106 | n/a | n/a | 76 | n/a | n/a |
| WT | 7863 | 92 | n/a | n/a | 20 | n/a | n/a |
| MM | 11083 | 21 | 13 | 8 | 19 | 17 | 2 |
| MM | 8884 | 145 | 133 | 12 | 43 | 40 | 3 |
| MM | 6389 | 65 | 50 | 15 | 2 | 1 | 1 |
| MM | 5084 | 34 | 6 | 28 | 27 | 12 | 15 |
| MM | 8572 | 90 | 78 | 12 | 41 | 37 | 4 |
| MM | 5967 | 231 | 199 | 32 | 185 | 170 | 15 |
