## Supplementary material for "Characterization of the intraspecies chimeric mouse brain at embryonic day 12.5": Figure S2

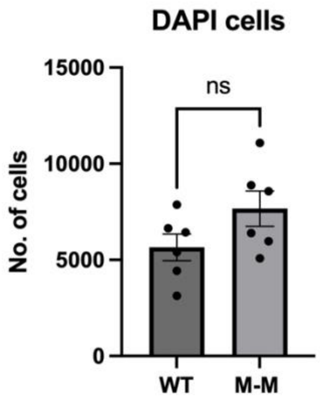


**Figure S2.** Total number of DAPI cells detected in the brain regions analyzed for microglia-like and macrophage-like cells (unpaired t-test, p=0.1105.
